# High-throughput peptide-centric local stability assay extends protein-ligand identification to membrane proteins, tissues, and bacteria

**DOI:** 10.1101/2025.04.28.650974

**Authors:** Kejia Li, Clement M Potel, Isabelle Becher, Nico Hüttmann, Martin Garrido-Rodriguez, Jennifer Schwarz, Mikhail M Savitski

## Abstract

Systematic mapping of protein-ligand interactions is essential for understanding biological processes and drug mechanisms. Peptide-centric local stability assay (PELSA) is a powerful tool for detecting these interactions and localizing potential binding sites. However, its original workflow is limited in throughput, sample compatibility and accessible protein targets. Here, we introduce a high-throughput adaptation - HT-PELSA - that increases sample processing efficiency 100-fold while maintaining high sensitivity and reproducibility. HT-PELSA substantially extends the capabilities of the original method by enabling sensitive protein-ligand profiling in crude cell, tissue and bacterial lysates, making it possible to identify membrane protein targets in diverse biological systems. We demonstrate that HT-PELSA can precisely and accurately determine binding affinities of small molecule inhibitors, sensitively detect direct and allosteric ATP binding, and reveal off-target interactions of a marketed kinase inhibitor in heart tissue. By enhancing scalability, reducing costs, and enabling system-wide drug screening across a wide range of sample types, HT-PELSA - when combined with next-generation mass spectrometry - offers a powerful platform poised to accelerate both drug discovery and basic biological research.

## Introduction

Protein function is often dynamically regulated by interactions with small molecules. Understanding the molecular mechanisms behind processes such as metabolic reactions and drug responses requires systematic techniques capable of probing protein-ligand interactions on a proteome-wide scale^1^. Among the various proteome-wide methods developed for mapping these interactions^2–5^, the peptide-centric local stability assay (PELSA)^6^ has recently emerged as a powerful tool due to its simplicity and high performance. This approach detects twice as many interactions as other techniques, while accurately pinpointing binding regions^6^. PELSA employs limited proteolysis^7^ to identify protein regions stabilized by ligand binding. Unlike earlier mass spectrometry-based approaches^2,4^, PELSA specifically analyzes peptides released from proteins after a brief trypsin digestion pulse, followed by the removal of undigested proteins using a molecular weight filter-based strategy.

The original PELSA workflow has two key limitations: it processes each sample individually, which restricts its throughput, and is performed on centrifuged lysates, confining the method primarily to cytoplasmic proteins. Here, we introduce an optimized, high-throughput adaptation, HT-PELSA, in which all steps are performed in 96-well plates at room temperature, improving throughput 100-fold. This streamlined workflow enables the preparation of hundreds of samples per day, enhancing scalability and reproducibility while reducing costs without compromising performance. Moreover, we demonstrate that HT-PELSA expands PELSA’s capabilities to measure membrane protein targets, across a broader range of sample types - including cell lines, tissues, and lower organisms. This advancement paves the way for more comprehensive profiling of protein-ligand interactions and offers powerful new opportunities for both drug discovery and fundamental biological research.

## Results

### High-throughput PELSA workflow

Compared to the original PELSA workflow, HT-PELSA introduces several key improvements that enhance efficiency, scalability, reproducibility and cost effectiveness (Figure 1A): (i) All steps are performed in 96-well plates instead of single tubes, substantially increasing throughput; (ii) All 96 samples are processed simultaneously, unlike the original PELSA protocol, where each sample was prepared sequentially due to short incubation times. This reduces processing time up to a 100-fold depending on the number of samples, enhances reproducibility, minimizes human error, and ensures uniform ligand incubation time across all samples; (iii) Samples are processed at room temperature, which eliminates the need for incubation at 37°C, streamlining the procedure (Figure S1A, Supplementary Dataset 1); (iv) The digestion time is extended to 4 minutes to ensure ease of operation without compromising performance (Figure 1B, Figure S1A, Supplementary Dataset 1); (v) Intact, undigested proteins are removed using 96-well C18 plates, which selectively retain large protein fragments while allowing shorter peptides to elute. This replaces molecular weight cut-off filter single units, thus increasing throughput and eliminating the need for additional desalting before mass spectrometry analysis. Moreover, this procedure is compatible with crude lysates cell lines, tissues, and bacteria - samples that would otherwise clog filter membranes.

**Figure 1:**
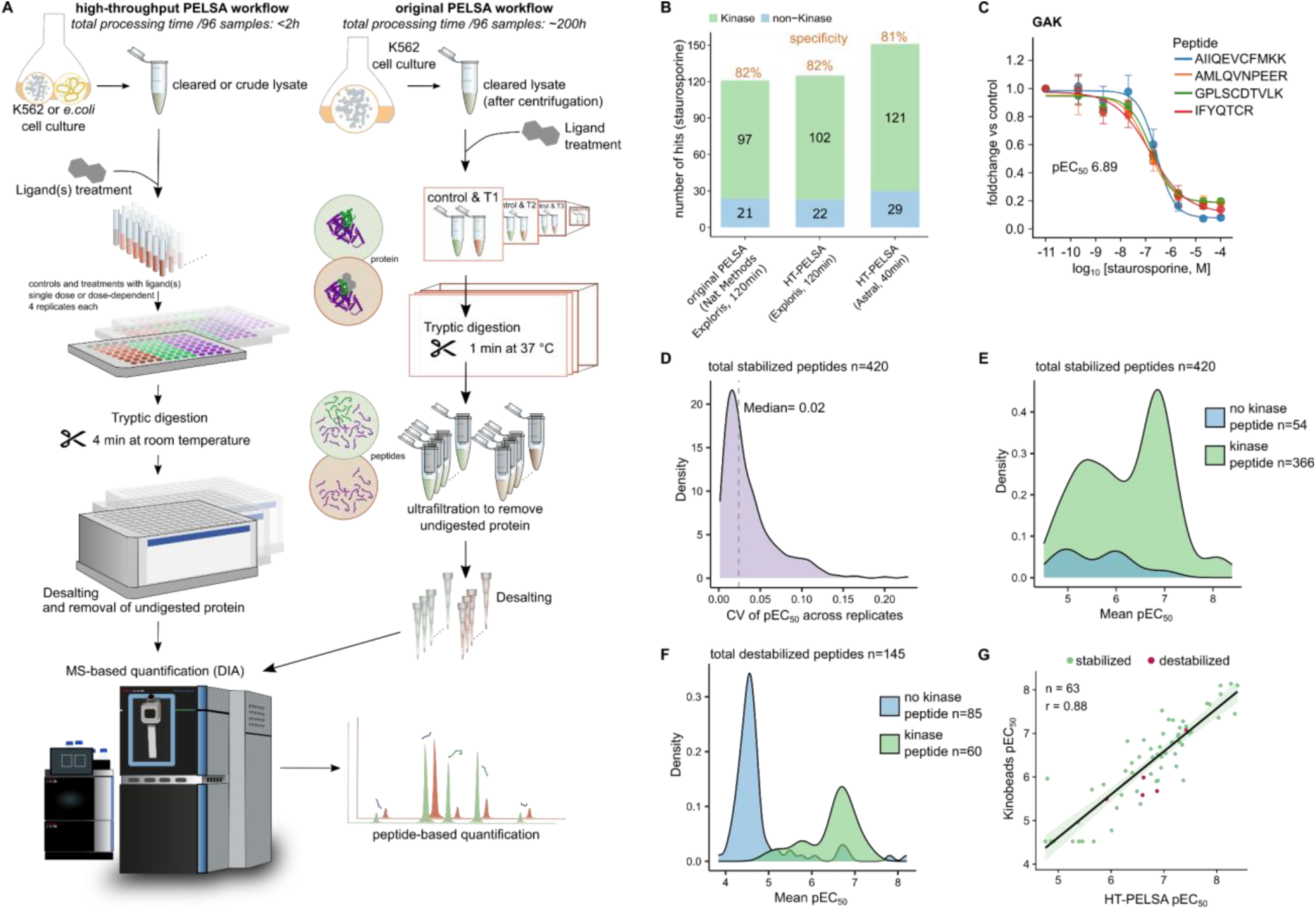
HT-PELSA enables precise and accurate identification of drug targets and their binding affinities (A) Comparison of the workflows between the high-throughput HT-PELSA platform and the original PELSA. In the high-throughput platform, each step is performed in a 96-well format. (B) The number of kinase and non-kinase targets identified in staurosporine binding experiments in the original PELSA dataset^6^ (Exploris), HT-PELSA (Exploris), and HT-PELSA (Astral) platforms are compared. All data were processed uniformly using DIA-NN (version 1.8.1). The specificity of kinase targets among all identified targets is also indicated. (C) The peptides of the Cyclin-G associated kinase (GAK) that exhibit a dose-dependent response to staurosporine (R² > 0.9, where R² denotes the correlation to a sigmoidal drug-response curve) are consistent across all four replicates. Error bars represent the standard deviation of fold change across the four PELSA replicates. These peptides, all derived from the kinase domain of GAK, show similar affinities for staurosporine, with pEC_50_ values of 6.59, 6.89, 6.77, and 6.81. (D) The coefficient of variation distribution of staurosporine pEC_50_ values for all stabilized peptides. Peptides were considered stabilized if they exhibited a dose-dependent response (R² > 0.9) and were at least 30% more resistant to trypsin at the highest staurosporine concentration, across a minimum of three PELSA replicates. (E) The staurosporine pEC_50_ distribution of all stabilized peptides, categorized by kinase versus non-kinase groups. For all dose-responsive peptides, pEC_50_ values represent the mean across replicates. (F) As in (E), but for all destabilized peptides. Destabilized peptides are defined as those with R² > 0.9 and at least 30% more susceptible to trypsin at the highest staurosporine concentration, across a minimum of three replicates. (G) Pearson correlation of staurosporine pEC_50_ values obtained from HT-PELSA and kinobeads assay. Each dot represents a kinase identified in both datasets, with the number of kinases labeled.

Overall, this workflow reduces sample processing time to under two hours for up to 96 samples - with the possibility to process multiple plates in parallel - from cell lysis to mass spectrometry-ready peptides. This offers a scalable, efficient and robust alternative to the original protocol. To validate the performance of HT-PELSA, we compared our high-throughput approach with the original method by identifying protein targets of staurosporine, a broad-spectrum kinase inhibitor commonly used as a benchmark for protein-ligand interaction studies. As shown in Figure 1B, the total number of kinases identified as binding to staurosporine, as well as the method specificity, is comparable to that of the original protocol when analyzed on the same instrument (Orbitrap Exploris), confirming the reliability and effectiveness of HT-PELSA. Notably, running the same samples on the new-generation mass spectrometer (Orbitrap Astral^8^) improves throughput 3-fold, and increases the number of identified targets by 22% (Figure 1B, Supplementary Dataset 1). 93% of the significantly stabilized kinase peptides were localized within or near kinase domains (Figure S1B,C, Supplementary Dataset 1), thus confirming that HT-PELSA accurately maps ligand binding regions.

### Systematic determination of protein-ligand binding affinity

In order to understand the biological or therapeutic implications of a protein-ligand interaction, it is essential not only to identify the interaction but also to characterize its binding affinity. Our high-throughput approach facilitates the reproducible measurement of protein-ligand interactions across different ligand concentrations, allowing us to generate dose-response curves on a proteome-wide scale and determine EC_50_ values (half-maximum effective concentration) for each protein target. These insights into binding dynamics are crucial for drug development, facilitating the identification and optimization of potential therapeutic compounds.

To validate our methodology, we first determined the pEC_50_ values for kinase-staurosporine interactions. HT-PELSA enables parallel processing of hundreds of samples, which results in highly reproducible pEC₅₀ values across replicates (median coefficient of variation = 2%), demonstrating its precision in quantifying ligand binding affinities (Figure 1C,D, Supplementary Dataset 2). Stabilized peptides - those less susceptible to tryptic digestion - showed high selectivity for kinases (90%; Figure 1E). Among the non-kinase stabilized peptides, several known off-targets of staurosporine were identified, such as Ferrochelatase (FECH)^3,9^ and OSBPL3^3^ (Supplementary Dataset 2), underscoring how dose–response analysis increases confidence in target identification. Notably, destabilized peptides contained a lower proportion of kinase-derived sequences (Figure 1F, Supplementary Dataset 2). However, those corresponding to kinases exhibited higher pEC₅₀ values compared to their non-kinase counterparts (Figure 1F, Supplementary Dataset 2). This suggests that while stabilized peptides are generally more reliable for target discovery, incorporating dose–response curves enhances both the specificity and confidence of hits identified through destabilization. Moreover, it supports the observation seen in thermal proteome profiling (TPP)^3^ that a small proportion of kinase targets consistently gets destabilized upon inhibitor binding.

Remarkably, the binding affinities of kinases measured by HT-PELSA closely aligned with values obtained from previously published kinobeads competition assays^10,11^ - the gold standard for systematic measurement of kinase-inhibitor affinities. This agreement demonstrates that, in addition to its high precision, HT-PELSA provides highly accurate affinity measurements for both stabilized and destabilized targets (Figure 1G, Supplementary Dataset 2).

Next, we applied HT-PELSA to profile the binding affinity of the metabolite ATP. Using a dose– response assay we identified a total of 1,426 stabilized peptides, enabling the characterization of ATP binding affinities for 301 *Escherichia coli* proteins (Figure 2A, B, Supplementary Dataset 3). A high proportion of stabilized peptides (1,013; 71%) and of stabilized proteins (174; 58%) corresponded to Uniprot-annotated ATP binders. This represents a substantial leap in coverage and specificity compared to the previous most comprehensive study systematically profiling protein-ATP interactions, using limited proteolysis^12^ (Figure 2C, Figure S2A, Supplementary Dataset 3). This enhanced performance was also evident when taking into account a single high ATP concentration. At 5 mM ATP, HT-PELSA detected 172 known ATP-binding proteins with 61% specificity, while LiP-MS detected 66 ATP binders (41% specificity) at the same concentration, and 84 (36% specificity) at 25 mM ATP^12^ (Figure S2B, Supplementary Dataset 3).

**Figure 2:**
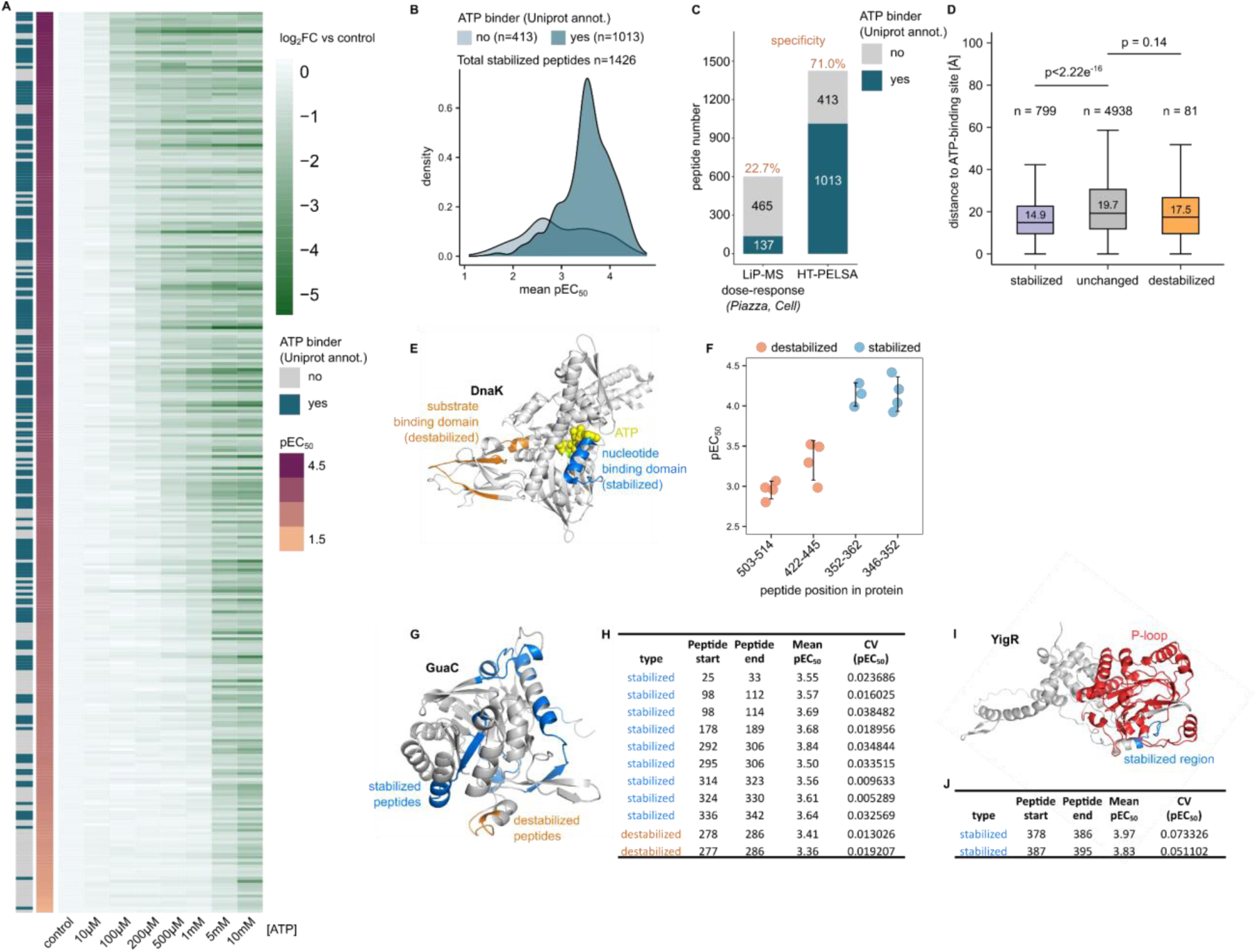
HT-PELSA enables high-resolution characterization of ATP binding proteins in *Escherichia coli* (A) Heatmap showing the log₂ fold change (relative to control) of 301 proteins exhibiting dose-dependent stabilization by ATP. Proteins are annotated based on whether they are UniProt-annotated ATP-binding proteins and are ranked by their apparent ATP-binding affinity (pEC_50_). (B) Distribution of pEC_50_ values for all peptides stabilized by ATP (defined as R² > 0.9 and ≥30% increased resistance to trypsin at the highest ATP concentration in at least three replicates), grouped by whether they originate from UniProt-annotated ATP-binding proteins. (C) Comparison of the number of peptides exhibiting dose-dependent responses to ATP in a published LiP-MS dataset^12^ and HT-PELSA, grouped by whether they are derived from UniProt-annotated ATP-binding proteins. (D) Distributions of the Euclidean distance between peptide cleavage sites and UniProt-annotated ATP-binding residues for peptides belonging to known ATP-binders, grouped by stabilized peptides (n = 799), unchanged peptides (n = 4,938), and destabilized peptides (n = 81). Statistical significance was assessed using a two-sided Wilcoxon rank-sum test (no adjustment). Box plots show the median (line), interquartile range (box), and ±1.5× IQR (whiskers); outliers are omitted for clarity. (E) Structural representation of DnaK (PDB: 4B9Q), with stabilized peptides highlighted in blue and destabilized peptides in orange. ATP is shown as yellow spheres. (F) pEC_50_ values of peptides from DnaK stabilized and destabilized by ATP across PELSA replicates. Error bars indicate standard deviations. (G) Structural model of GuaC (AlphaFold: AF-P60560), with nine peptides stabilized by ATP shown in blue and two destabilized peptides in orange. (H) Summary of mean pEC_50_ values and coefficients of variation across PELSA replicates for peptides displayed in (G). (I) Structural model of YjgR (AlphaFold: AF-P39342), with the P-loop NTPase domain colored in red and two peptides stabilized by ATP shown in blue. (J) Summary of mean pEC_50_ values and coefficients of variation across PELSA replicates for peptides shown in (I).

The comprehensive, dose-dependent measurements with HT-PELSA also allow for the precise determination of pEC₅₀ values for the ATP-stabilized proteins identified (Figure 2B, Figure S2C, Supplementary Dataset 3). Similar to the case of staurosporine and kinases, a smaller proportion of destabilized peptides corresponds to annotated ATP-binding proteins and exhibits, on average, higher pEC₅₀ values (Figure S2D, S2E, Supplementary Dataset 3). Analysis of AlphaFold^13^ structures for proteins with known ATP binding sites revealed that stabilized peptides are significantly closer to these binding sites than peptides with no detectable changes (Figure 2D). Interestingly, this trend is much less pronounced - and not statistically significant - for destabilized peptides (Figure 2D), suggesting that these could reveal allosteric effects induced by ATP binding in distal regions of proteins. Consistently, the destabilized proteins are enriched in transporter complex or ATP-dependent membrane complex and chaperone proteins (Figure S2F, S2G, Supplementary Dataset 3). For the HSP70-like chaperone DnaK, the protein-folding activity depends on its ATP-regulated interaction with substrates, mediated by allosteric communication between the nucleotide-binding domain and the substrate-binding domain, with substrate release triggered by ATP binding^14^. We observe that DnaK is stabilized with high affinity at the ATP binding site, and destabilized with lower affinity in the substrate binding region (Figure 2E and 2F, Supplementary Dataset 3). This supports the notion that substrate release is triggered by ATP binding at sufficiently high concentrations, making the substrate-binding domain more accessible to trypsin. Among the non-annotated ATP binders, we identify GuaC, which shows both stabilization and destabilization in two distinct regions at nearly identical ATP concentrations (Figure 2G and 2H, Supplementary Dataset 3). In *Trypanosoma brucei*, GuaC has been shown to be inhibited by ATP through allosteric regulation^15^, and similar inhibition of GuaC activity by ATP has been previously reported in *E. coli*^16^. Finally, we identify YjgR, an uncharacterized protein with a P-loop NTPase domain, which is strongly stabilized by ATP. Notably, the stabilized peptides are located within or adjacent to the P-loop region (Figure 2I and 2J, Supplementary Dataset 3). In summary, our approach provides precise determination of ATP-binding affinities for both known and previously unannotated ATP-binding proteins. Furthermore, the high sequence-level resolution of our data not only detects binding events at ATP-binding sites but also offers valuable insights into allosteric regulation.

### Identification of membrane protein targets

Membrane proteins serve as the initial mediators of signaling cascades, making them prime targets for drug development. Originally, PELSA was performed on cleared lysates to avoid clogging of the mass cut-off filter, meaning these proteins were mostly removed by centrifugation before the ligand-binding assay. Here, we demonstrate that the plate-based HT-PELSA can be performed directly on crude lysates, preserving membrane proteins and expanding the assay’s applicability (Figure 3A).

**Figure 3:**
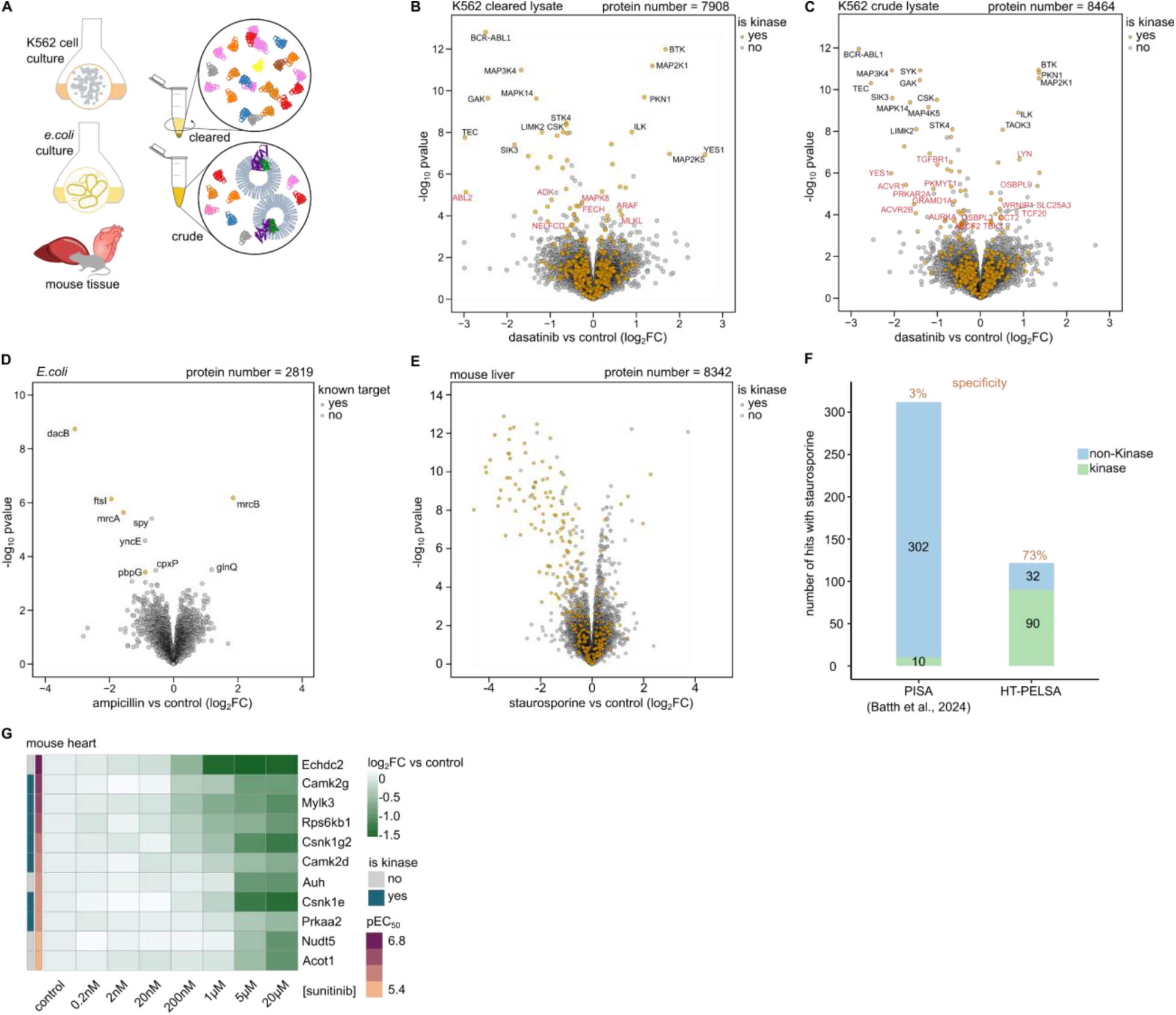
HT-PELSA extends target identification to membrane proteins in cell lines, mouse tissues and bacteria (A) Schematic representation of crude or cleared K562 cell, *E. coli*, and tissue lysates for HT-PELSA analysis. (B) Volcano plot showing candidate dasatinib-binding targets identified by HT-PELSA in cleared K562 lysates treated with 5 μM dasatinib. Proteins were considered candidate targets if they met the threshold of -log_10_P > 4. For each protein, the peptide with the lowest p-value is displayed. (C) As in (B), but for crude K562 lysates. Protein targets that are uniquely identified in the cleared lysates or crude lysates are annotated with gene names in red. (D) Volcano plot showing candidate ampicillin-binding targets identified by HT-PELSA in crude *E. coli* lysates exposed to 10 μM ampicillin. Proteins with -log_10_P > 3.4 are annotated with gene names. (E) Volcano plot showing candidate staurosporine-binding targets identified by HT-PELSA in crude mouse liver lysates exposed to 20 μM staurosporine. (F) Comparison of the performance of PISA^19^ and HT-PELSA in identifying staurosporine targets in mouse liver lysates. (G) Heatmap displaying log_2_ fold change (versus control) of 11 proteins which show dose-dependent stabilization to sunitinib in mouse heart lysate. The gene names of the proteins are labeled.

As proof of concept, we first identified targets of dasatinib, a clinically approved tyrosine kinase inhibitor used to treat certain types of leukemia. While its primary target BCR-ABL1 is an intracellular kinase, many of dasatinib’s off-targets are membrane-associated proteins^17^. HT-PELSA performed on crude K562 cell lysates reveals 556 additional proteins compared to cleared lysates (Figure 3B,C, Supplementary Dataset 4). These proteins are significantly enriched for Gene Ontology terms related to mitochondrial and plasma membrane functions (Figure S3A). Our results showed that the primary target BCR-ABL1 stands out as the top target for both cleared and crude lysates (Figure 3B,C, Supplementary Dataset 4). Beyond BCR-ABL1, we identified 43 shared targets in both crude and cleared lysates, 36 of which are protein kinases, highlighting the promiscuity of dasatinib (Figure S3B). However, among these 43 targets, only 3 are transmembrane proteins (Figure S3C). Performing HT-PELSA on crude lysates resulted in an increase in identification of membrane-associated target proteins, including several known dasatinib off-targets such as LYN, TGFBR1, YES1, ACVR1 and ACVR2B^17^ (Figure 3C, Figure S3B,C, Supplementary Dataset 4).

To further emphasize the ability to characterize membrane protein interactions, we next sought to identify ampicillin targets in *Escherichia coli* crude lysate. Ampicillin is a penicillin derivative antibiotic binding to inner membrane penicillin-binding proteins, which are essential for cell membrane biosynthesis. HT-PELSA identified 5 known targets of ampicillin (*mrcA*, *mrcB*, *ftsI*, *dacB* and *pbpG*, Figure 3D, Supplementary Dataset 5). In comparison, TPP in *E. coli* lysate identified 2 known targets, while TPP in live cells identified 4 known targets^18^. These findings also demonstrate that the HT-PELSA protocol effectively captures protein-ligand interactions in crude lysates of bacterial samples, expanding its applicability to microbial systems.

### HT-PELSA in mouse tissue reveals kinase inhibitors off-targets

Mapping protein-ligand interactions in tissue provides more biologically relevant insights than in cell lines. However, studying these interactions is more challenging due to heightened complexity and potential biomolecular contaminants, which can affect both the number and specificity of target identifications. For example, a recent study detected only a handful of kinases binding to staurosporine in mouse liver tissue^19^. In contrast, HT-PELSA successfully identifies 90 kinases in a mouse liver crude lysate, demonstrating comparable performance in both cell lines and tissues (Figure 3E,F, Supplementary Dataset 6).

Identifying drug targets and their affinities beyond the primary target is crucial in drug discovery as off-target interactions can result in harmful side effects. To illustrate this, we characterized the targets of sunitinib, a clinically approved multi-targeted tyrosine kinase inhibitor used to treat renal, gastrointestinal, and pancreatic cancers. Clinical studies have reported direct cardiomyocyte toxicity associated with sunitinib treatment, which can progress to heart failure in affected patients^20^. In this study, we identified four known kinase off-targets of sunitinib in mouse heart tissue, the α subunit Prkaa2 of Ampk, Camk2g, Csn1ke and Csnk1g2^17^ (Figure 3G, Supplementary Dataset 7). Inhibition of Ampk in the heart has been recognized as a key driver of sunitinib-induced cardiotoxicity by reducing mitochondrial ATP production^21^. We also characterized additional direct high-affinity off-targets of sunitinib, (Figure 3G). These include Camk2d, a kinase with heart specific expression^22^ that has so far only been reported to bind sunitinib in a recombinant kinase assay^23^ and is involved in regulation of heart function^24^ as well as the myosin light chain kinase 3 (Mylk3) that is known to have a cardiomyocyte specific expression^25^ underscoring the importance of tissue-specific drug target profiling. Mylk3 activity is essential for maintaining basal contractile function in the heart^26,27^, potentially revealing additional mechanisms underlying sunitinib-induced cardiotoxicity. In addition, several mitochondrial proteins (Echdc2, Auh, Acot1) were identified as direct high affinity targets of sunitinib (Figure 3G). Since mitochondrial functions are essential for energy homeostasis, their disruption may contribute to mitochondrial dysfunction and, ultimately, cardiac pathologies.

## Discussion

The recently introduced PELSA workflow enables sensitive and systematic probing of protein-ligand interactions. In this study, we develop HT-PELSA, a workflow that dramatically increases sample throughput and streamlines sample preparation. These advancements pave the way for high-throughput screening of protein–small molecule interactions, providing information on protein target, binding affinity and protein binding regions. Furthermore, the introduced HT-PELSA protocol extends protein-ligand profiling beyond the cleared cell line lysate setting. We demonstrated that HT-PELSA accommodates a broad range of sample types, including crude tissue and bacterial lysates, and enables mapping of ligand interactions with membrane proteins, a key class of proteins in drug discovery.

HT-PELSA unlocks exciting new possibilities. The 96 well plate format, extended digestion time of four minutes and removal of the short 37°C incubation make the workflow primed for future automation. With higher sensitivity than alternative approaches - currently requiring 60 μg of protein input per sample, with only 5% of the resulting peptides being injected for mass spectrometry analysis - the input could be further scaled down. This makes HT-PELSA particularly advantageous for studying protein interactions in limited or valuable materials, such as clinical samples. Finally, in the future, the scope of HT-PELSA could be expanded to systematically investigate protein-protein or protein–nucleic acid interactions, further broadening its impact in biomolecular research.

## Data availability

All results and data are available online (https://2u7b8b-nico0h0ttmann.shinyapps.io/PELSAAPP/). All raw files, search parameters and search outputs were deposited to the ProteomeXchange Consortium through the PRIDE partner repository with the dataset identifier PXD062869.

## Materials and Methods

### Human Cell lines

K562 cells (ATCC CCL-243) were cultured in RPMI 1640 medium (Thermo Fisher) supplemented with 2 mM L-glutamine, and 10% FBS at 37°C (5% CO_2_). For collection, K562 cells were pelleted at 1,000 x g for 5 min. The collected cells were washed three times with ice-cold PBS and stored at −80 °C for further analysis.

### Escherichia coli

(A) *E. coli* K-12 strain BW25113 cells were grown overnight at 37°C in lysogeny broth (LB Lennox) and diluted 100-fold into 20 mL of fresh LB. Cultures were grown aerobically at 37°C with shaking until optical density at 578 nm (OD578) of 2. For collection, *E. coli* were pelleted at 1,000 x g for 5 min and washed three times with ice-cold PBS and stored at −80 °C for further analysis.

### Mouse Tissues

All mouse experiments were performed using approved protocols by the EMBL ethics committee (license 21-002_HD_MZ). For the liver collection: germ-free C57BL/6 mice were maintained and bred in gnotobiotic isolators (CbC) with a 12-h light–dark cycle. Mice were provided with standard, autoclaved chow (1318P FORTI, Altromin) ad libitum. After reaching maturity, at 7 weeks, a male mouse was colonized with laboratory strains from a cryopreserved inoculum composed of *A. muciniphila* DSM22959, *B. uniformis* DSM6597, *C. sporogenes* ATCC15579, *E. rectale* DSM17629, *L. gasseri* DSM20243, *P. distasonis* DSM20701, *S. copri* DSM18205, and *R. gnavus* ATCC29149. The colonized animal was single-housed for 14 days before it was killed and liver samples were collected and stored at −70 °C until further analysis.

For the mouse heart collection: C57BL/6J mice were maintained in individually ventilated plastic cages (Tecniplast) under a 12-hour light/dark cycle, with ad libitum access to 1318 P autoclavable diet (Altromin, Germany). At four months of age, a male mouse was killed and heart samples were collected and stored at −70 °C until further analysis.

### Lysate preparation

K562 cells were resuspended in ice-cold lysis buffer (PBS supplemented with 1% (v/v) protease inhibitor cocktail, Sigma P8340-5ML) and lysed by three cycles of snap-freezing in liquid nitrogen followed by thawing in a 37 °C water bath. To obtain cleared lysates, cell extracts were centrifuged at 20,000 × g for 10 minutes at 4 °C to remove cellular debris. For crude lysates, non-centrifuged cell extracts were used directly for subsequent analysis.

*E.coli* cells were resuspended in lysis buffer containing PBS, 1% (v/v) protease inhibitor cocktail, 50 µg/mL lysozyme (Sigma, L7386), 0.25% (v/v) benzonase (Merck, 71206-3), and 2 mM MgCl₂. The suspension was incubated at 21 °C for 20 minutes to facilitate cell wall degradation. Following incubation, cells were further lysed by three cycles of freeze-thaw. For ampicillin HT-PELSA analysis, the resulting crude *E. coli* lysates were used directly. For ATP HT-PELSA analysis, the lysis buffer was supplemented with an additional 10 mM MgCl₂ to promote ATP-Mg²⁺ complex formation at the maximum working ATP concentration of 10 mM. To enable accurate determination of the binding affinity to ATP, lysates were desalted using size-exclusion chromatography (SEC, Zeba Spin Desalting Columns 7 kDa molecular weight cut-off, Thermo Fisher Scientific) to remove endogenous ATP. Before desalting, the crude *E. coli* lysates were spun down at 500 x g, 4 °C for 10 min to remove intact cells which could potentially block the SEC columns. The resulting desalted *E. coli* lysates were used for subsequent analysis.

The mouse liver tissue was resuspended in lysis buffer containing PBS and 1% (v/v) protease inhibitor cocktail, then homogenized using a bead beater (Beadruptor Elite, Omni International) with a mix of 2.8 mm and 1.6 mm zirconium silicate beads for three cycles of 20 s at 4 m/s, with 30 s cooling between cycles. The resulting lysates were used for subsequent analysis. Mouse heart lysates were prepared similarly to liver lysates, but with five cycles of bead beating for 20 seconds at 6 m/s, with 30-second cooling intervals between cycles.

For all samples, the protein concentrations of lysates were measured with Rapid Gold BCA protein Assay Kit (Thermo Scientific) and adjusted to 1.2 mg/mL with the lysis buffer.

### Incubation with the ligands

In benchmarking experiments against the original PELSA protocol^6^, a fixed staurosporine concentration of 20 μM was used for both HT-PELSA protocols (4-min and 3-min digestion at room temperature). For the staurosporine dose-response assay, concentrations tested were 0.2 nM, 2 nM, 20 nM, 200 nM, 2 μM, 20 μM, and 100 μM. In the *E. coli* ATP dose response experiment, ATP concentrations of 10 μM, 100 μM, 200 μM, 500 μM, 1 mM, 5 mM, and 10 mM were used. In the dasatinib experiment, both crude and cleared K562 cell lysates were incubated with 5 μM dasatinib. In the *E. coli* ampicillin binding assay, crude *E. coli* lysates were incubated with 10 μM ampicillin. In the mouse liver staurosporine binding experiment, lysates were incubated with 20 μM staurosporine. In the mouse heart sunitinib dose response experiment, concentrations of 0.2 nM, 2 nM, 20 nM, 200 nM, 1 μM, 5 μM, and 20 μM were used. Stock solutions (100-fold) of staurosporine, dasatinib, and sunitinib were prepared in dimethyl sulfoxide (DMSO), with equivalent DMSO volumes added to the control group. Stock solutions (50-fold) of ATP and ampicillin were prepared in water, with equivalent water volumes added to the control group. The pH of the ATP stock solution was adjusted with sodium hydroxide to 7 prior to use. The lysates were incubated with the different ligands at 25 °C for 30 min.

### HT-PELSA workflow

After incubation, each sample was divided into four 50 μL replicates in a 96-well plate. Using a Gilson Platemaster, all samples were transferred to a second 96-well plate pre-loaded with 5 μL of trypsin (Sigma, cat. no. T1426) stock solution (5 mg/mL). The trypsin was kept on ice and added to the plate shortly before sample addition. The samples were mixed with trypsin by pipetting up and down for 30 s and incubated at room temperature for an additional 3 min and 30s. Digestion was quenched by adding a threefold volume (165 μL) of 8 M guanidine hydrochloride (GdmCl) solution (containing 8 M GdmCl and 60 mM HEPES, pH 8.2), resulting in a final GdmCl concentration of 6 M. Control samples were subsequently supplemented with the same concentration of ligand as their corresponding ligand-treated samples, to account for potential interference with ionization during mass spectrometry analysis. 10 mM of Tris(2-carboxyethyl)phosphine (TCEP) and 40 mM of chloroacetamide (final concentrations) were added to the samples and the mixture was then heated at 95 °C for 5 min for cysteine carbamidomethylation. Alternatively, reduction/alkylation can be performed at lower temperature and longer incubation time.

After carbamidomethylation, the samples were acidified with trifluoroacetic acid (TFA) to a final concentration of 1%. Prior to loading onto the 100 mg SepPak tC18 96-well plate, the plate was conditioned as follows, with every step being performed at room temperature: first, it was centrifuged at 3,000 × g for 5 minutes without liquid to account for potential differences in C18 packing that could affect flow rate. Each well was then washed twice with 1 mL of 100% ACN (centrifuged at 20 × g for 1 minute), followed by two washes with 1 mL of 0.1% TFA (centrifuged at 100 × g for 1 minute with an acceleration setting of 5 out of 9). Samples were then loaded onto the plate (centrifuged at 100 × g for 1 minute with acceleration set to 5/9) and washed twice with 1 mL of 0.1% TFA (centrifuged as before). Finally, peptides were eluted with 2 × 100 µL of 50% ACN, 0.1% TFA (centrifuged at 20 × g for 1 minute), followed by a final centrifugation step at 500 × g for 1 minute. The eluted peptides were collected directly onto a mass spectrometry-compatible 96-well plate and dried. To ensure that the tips of the C18 plate does not come into contact with the bottom of the well, a 96-tip holder from a pipette tip box was used in between the C18 plate and the collection plate. Alternatively, only a fraction of the elution can be dried, as only 5% of the eluate was injected into the LC-MS/MS system.

### LC-MS/MS Analysis

The samples were resuspended in a loading buffer containing 0.1% TFA, 4% acetonitrile in MS-grade water and a volume corresponding to 5% of the sample was injected. For both LC systems, Solvent A was 0.1% formic acid supplemented with 3% DMSO in LC–MS-grade water and solvent B was 0.1% formic acid supplemented with 3% DMSO in LC–MS-grade acetonitrile. For the HT-PELSA Exploris 480 analysis, peptides were separated using an UltiMate 3000 RSLCnano system (Thermo Fisher Scientific) equipped with a trapping cartridge (Precolumn; C18 PepMap 100, 5 μm, 300-μm inner diameter × 5 mm, 100 Å) and an analytical column (Waters nanoEase HSS C18 T3, 75 μm × 25 cm, 1.8 μm, 100 Å). Peptides were loaded onto the trapping cartridge (30 μl/min solvent A for 3 min) and eluted with a constant flow of 300 nL/min. Peptides were separated using a linear gradient of 8–25% B for 99 min, followed by an increase to 40% B within 5 min before washing at 85% B for 4 min and re-equilibration to initial conditions. The LC system was coupled to an Exploris 480 mass spectrometer (Thermo Fisher Scientific) operated in data independent acquisition mode. The instrument was operated in positive ion mode with a spray voltage of 2.5 kV and a capillary temperature of 275 °C. Full-scan MS spectra were acquired with a resolution of 60,000 with a mass range of 420-680 and AGC of 3e^6^. DIA spectra were acquired in the Orbitrap mass analyzer with 4 m/z windows between 430 and 670 m/z. MS2 scan range was set to 200-1800 m/z, the normalized collision energy to 28 and the default charge state to 2+. The normalized AGC target was set to 3000% and the maximum injection time to auto.

For HT-PELSA Astral analysis, peptides were separated using an Vanquish Neo UHPLC system (Thermo Fisher Scientific) operated in trap-and-elute mode. The LC was equipped with a trapping cartridge (Precolumn; PepMap Neo C18, 5 μm, 300-μm inner diameter × 5 mm, 100 Å) and an analytical column (Ionopticks AUR3-25075C18-XT, 25 cm x 75 μm ID, 1.7 μm C18). Solvent A was 0.1% formic acid supplemented with 3% DMSO in LC–MS-grade water and solvent B was 0.1% formic acid supplemented with 3% DMSO in LC–MS-grade acetonitrile. Peptides were loaded onto the trapping cartridge and separated on the analytical column using a linear gradient of 5–26% B (flow rate of 300 nL/min) for 29.7 min, followed by an increase to 40% B within 3 min before washing at 85% B for 5 min and re-equilibration to initial conditions (total MS acquisition time of 40 minutes). The LC system was coupled to an Orbitrap Astral mass spectrometer (Thermo Fisher Scientific) operated in data independent acquisition mode. The instrument was operated in positive ion mode with a spray voltage of 1.8 kV and a capillary temperature of 280 °C. Full-scan MS spectra were acquired in the Orbitrap at a resolution of 240,000 with a mass range of 430-680 m/z, an AGC target of 5e^6^ charges and a maximum injection time of 5 ms. DIA spectra were acquired in the Astral mass analyzer with 2 m/z windows between 430 and 680 m/z. MS2 scan range was set to 150-2000 m/z, the normalized collision energy to 25 and the default charge state to 2+. The normalized AGC target was set to 500% and the maximum injection time to 5 ms.

### Data analysis

Raw files were analyzed using DIA-NN 1.8.1^28^ with directDIA module using an in silico DIA-NN predicted spectral library (C carbamidomethylation and N-terminal M excision; 2 missed cleavage; precursor m/z range: 430-680). The spectral library was generated from fasta files downloaded from Uniprot (Human: UniProt 2022 release, 20,311 sequences, reviewed; Mouse: Uniprot 2023 release, 17,173 sequences, reviewed; *E. coli*: Uniprot 2024 release, 4,531 sequences, reviewed). The DIA-NN search used the following parameters: Precursor FDR (%)= 1; Mass accuracy, MS1 accuracy, and Scan window = 0; Use isotopologues; MBR enabled; Heuristic protein inference enabled; No shared spectra; Protein inference = Genes, Neural network classifier: Single-pass mode; Quantification strategy: Robust LC (high precision); Cross-run normalization: RT-dependent; Library generation: Smart profiling; Speed and RAM usage: optimal results.

The precursor matrix outputs from DIA-NN were used for downstream statistical analysis in R. Precursor intensities were aggregated for each peptide to obtain peptide-level intensities. Peptides with any missing values were removed and the remaining peptides were subjected to empirical Bayes moderated t-statistics analysis between ligand-treated samples and control samples. To enable protein-level screening, each protein was represented by the peptide with minimal P value (two-sided empirical Bayes t-test) among all its quantified peptides, that is, representative peptides. The protein-level volcano plot was generated by using −log_10_P and log_2_FC of the representative peptide as the y and x axis, respectively. The empirical Bayes t-test P value is used to rank the probability of a protein being recognized as a target hit. Unless otherwise stated, only proteins displaying an increased stability by ligand treatment (that is, log2FC < 0) are considered for candidate target protein ranking.

For dose-response data analysis, the peptide-level intensities were calculated based on their respective precursors as above. Peptides are required to be quantified in at least three replicates of the dose-response experiment. For each replicate, the ratio (treated vs. control) was calculated by dividing the peptide intensity at each ligand concentration by the control group’s intensity. Prior to fitting dose-response curves, peptides were pre-filtered based on a minimum 30% stabilization or destabilization at the highest ligand concentration. Additionally, for stabilized peptides, the ratio at the second-highest concentration was required to be below 1; for destabilized peptides, it was required to be above 1. The ratio of peptide intensity at each ligand concentration was fitted to the ligand concentrations using a four-parameter log-logistic model in R (drc package). The -log_10_ transformed half-effective concentration (pEC_50_) was extracted from the fitting model. The correlation (R^2^) to a sigmoid trend of the ligand dose–response profile for each peptide was derived by performing a Pearson correlation analysis between the estimated values and the original values. For candidate target peptide determination, the R^2^ of peptides are required to be above 0.9 in at least three replicates; the mean value of the pEC_50_ values across different replicates as well as the coefficient of variation (CV) of the pEC_50_ values were calculated; peptides with CV of pEC_50_ > 0.25 are excluded. The peptide with the highest pEC_50_ was assigned to the corresponding protein to obtain protein-level pEC_50_ for each protein.

### Structural analysis

To evaluate the spatial relationships between peptides and annotated ligand-binding sites within proteins, we retrieved UniProt metadata files (.json) and AlphaFold-predicted protein structures (.pdb) for each unique protein of interest. Binding site annotations were extracted from the UniProt feature table by selecting entries labeled as “Binding site,” and the central residue of each annotated region was used as its representative. Protein structures were parsed using Biopython’s PDBParser, and per-residue AlphaFold confidence scores (pLDDT) were obtained from the B-factors of Cα atoms. For each peptide mapped to a protein, we calculated the minimum Euclidean distance between the Cα atom of the binding site residue and all atoms of the peptide’s start and end residues. The smaller of these distances is used as the distance between peptide and the binding site. These distances reflect straight-line measurements in three-dimensional space based on the static atomic coordinates provided by the AlphaFold-predicted models.

### Source of the known targets databases

The list of 523 human protein kinases was obtained from KinHub (http://www.kinhub.org). Kinase domain information, mouse kinase list, human trans-membrane protein list, and *E. coli* known ATP-binding proteins were all derived from UniProt. All lists can be found in the Supplementary Dataset 8.

### Gene ontology (GO) analysis

GO analysis for human cell lines and E. coli was performed with the *clusterProfiler*^29^ R package using the enrichGO and enricher functions. For GO analysis, the background was set as all quantified proteins in each dataset.

---

**Figure S1:**
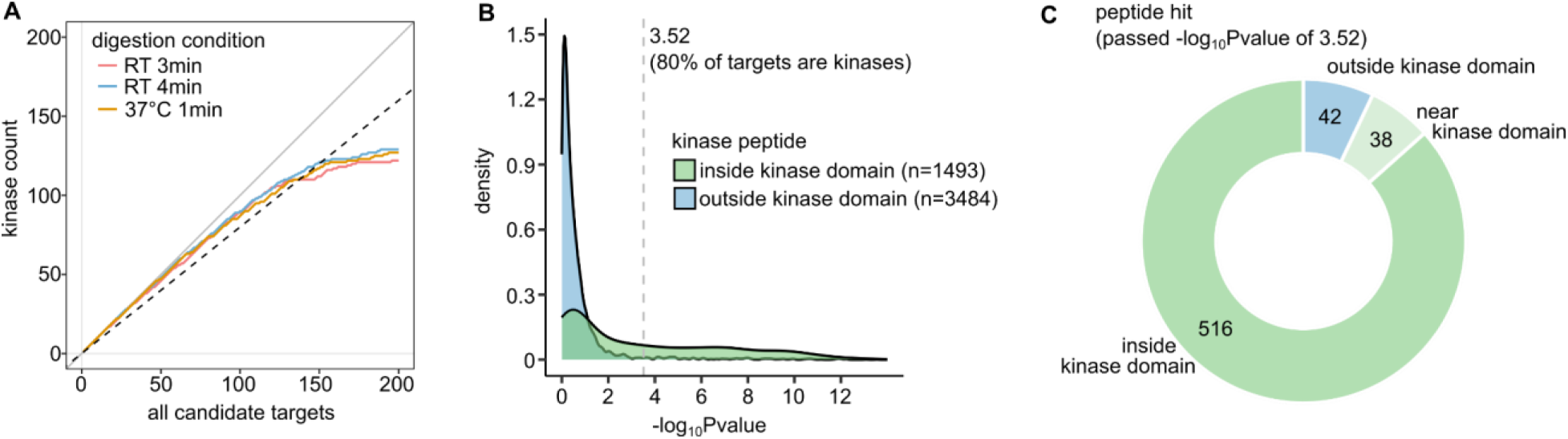
(A) True positive rate evaluation for different digestion conditions in HT-PELSA for staurosporine target identification. The gray line (slope = 1) and black dashed line (slope = 0.8) represent scenarios where 100% and 80% of the candidate targets are kinase targets, respectively. (B) Density plots showing −log_10_P-value distributions of all quantified peptides belonging to kinases, with tryptic cleavage sites located within or outside the kinase domains in HT-PELSA (RT 4 min) staurosporine analyses. The dashed line indicates the -log_10_P above which more than 80% of stabilized proteins are kinases. (C) The doughnut charts show the locations of kinase peptides meeting the −log_10_P cutoff from (B) and a log_2_FC (treated vs. control) < 0. “inside” denotes peptides with N- or C-termini within the kinase domain; “near” indicates N- or C-termini within 10 amino acids of the kinase domain; otherwise, peptides are classified as “outside”.

**Figure S2:**
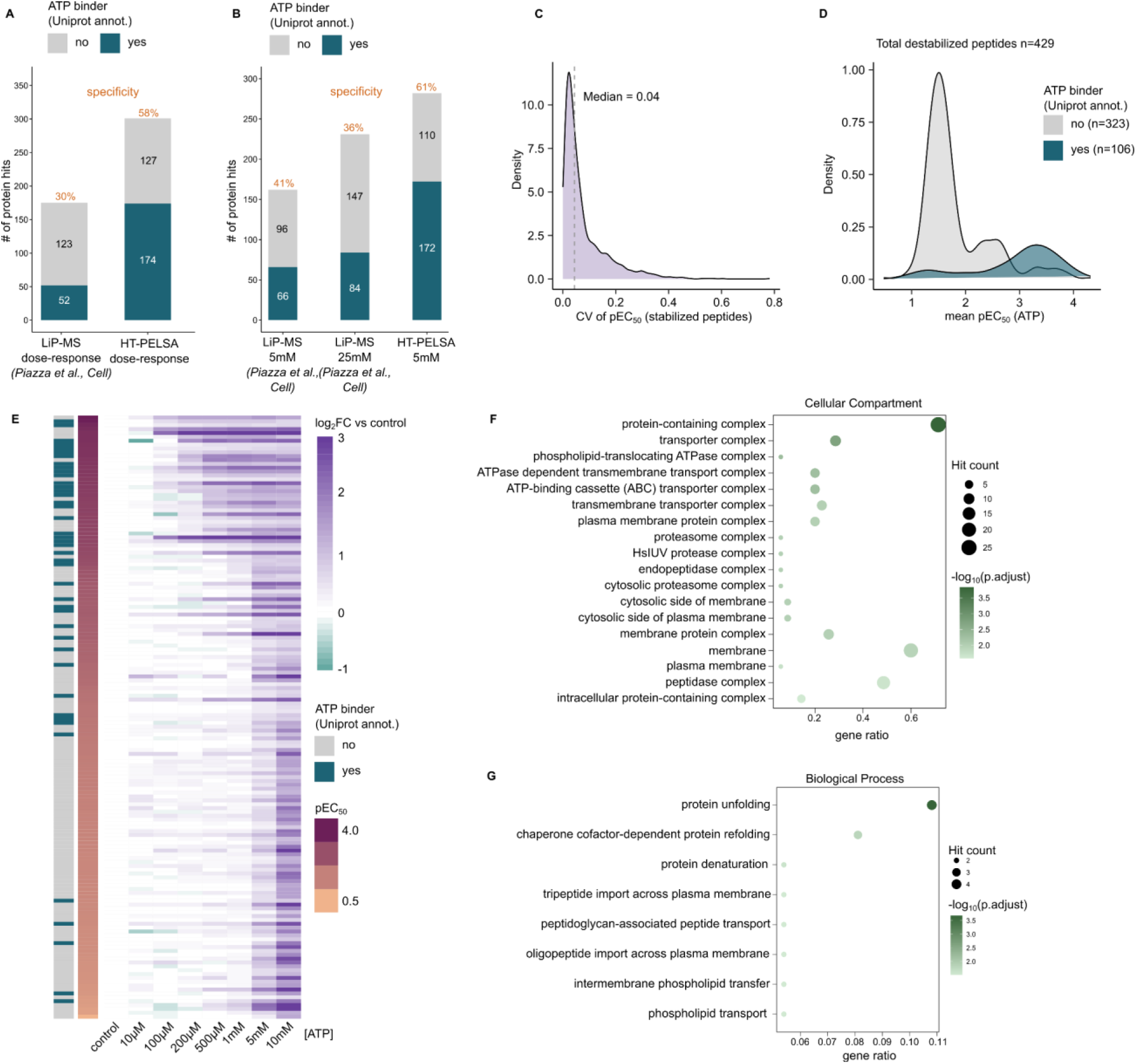
(A) Comparison of the total number of identified protein targets and Uniprot-annotated ATP binders in a published LiP-MS dataset^12^ and HT-PELSA based on their respective ATP dose-response data. (B) Comparison of the total number of identified protein targets and UniProt-annotated ATP-binding proteins at single ATP concentrations: 5 mM and 25 mM in LiP-MS^12^, and 5 mM in PELSA. (C) Distribution of the coefficient of variation of pEC_50_ values across PELSA replicates for all peptides exhibiting dose-dependent stabilization by ATP in HT-PELSA. Peptides were considered dose-dependently stabilized if they showed at least a 30% increase in resistance towards trypsin at the highest ATP concentration, with an R² > 0.9 in at least three replicates. (D) pEC_50_ distributions of all peptides exhibiting dose-dependent destabilization by ATP in HT-PELSA, grouped by whether they originate from UniProt-annotated ATP-binding proteins or not. Peptides were considered dose-dependently destabilized if they showed at least a 30% decrease in resistance towards trypsin at the highest ATP concentration, with an R² > 0.9 in at least three replicates. (E) Heatmap showing the log₂ fold change (relative to control) of 156 proteins exhibiting dose-dependent destabilization by ATP. Proteins are annotated according to whether they are UniProt-annotated ATP-binding proteins. (F) GO cellular component analysis of the high-affinity (pEC_50_ > 3) ATP-destabilized proteins (39 proteins total). (G) GO biological process analysis of the high-affinity (pEC_50_ >3) ATP-destabilized proteins (39 proteins total).

**Figure S3:**
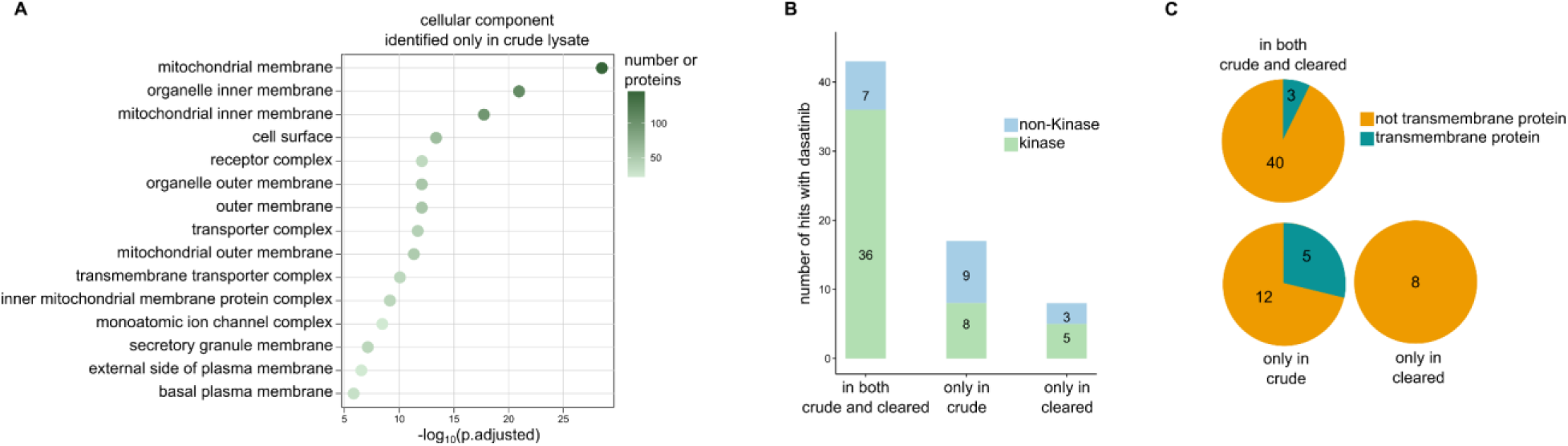
(A) Gene ontology (cellular component) analysis of 556 proteins that are uniquely identified in crude lysates versus cleared lysates of K562 cells. **(B)** Comparison of the numbers of candidate dasatinib-binding targets identified in crude and cleared K562 cell lysates, from left to right: targets identified in both crude and cleared lysates, only in the crude lysates, and only in the cleared lysates. **(C)** The percentage of trans-membrane targets for each category defined in (B).

## Acknowledgements

We thank Amber Brauer from the Zimmermann lab, Ernesto de la Cueva Bueno, and Alessandro Grassi from the Laboratory Animal Resources (LAR) at EMBL for providing the mouse tissues. We thank Suparat Scheu from the Proteomics Core Facility at the EMBL for providing the *E. coli* pellets. We thank Mira Lea Burtscher, Dimitrios Papagiannidis, and other members from Savitski lab for insightful discussions. This work was supported by the EMBL. M.G.-R. is supported through state funds approved by the State Parliament of Baden-Württemberg for the Innovation Campus Health + Life Science Alliance Heidelberg Mannheim. M.M.S. is supported by the Allen Distinguished Investigator award through the Paul G. Allen Frontiers Group.

