## Supplementary figures and images for "High-throughput peptide-centric local stability assay extends protein-ligand identification to membrane proteins, tissues, and bacteria"

### Supplementary Figure 1

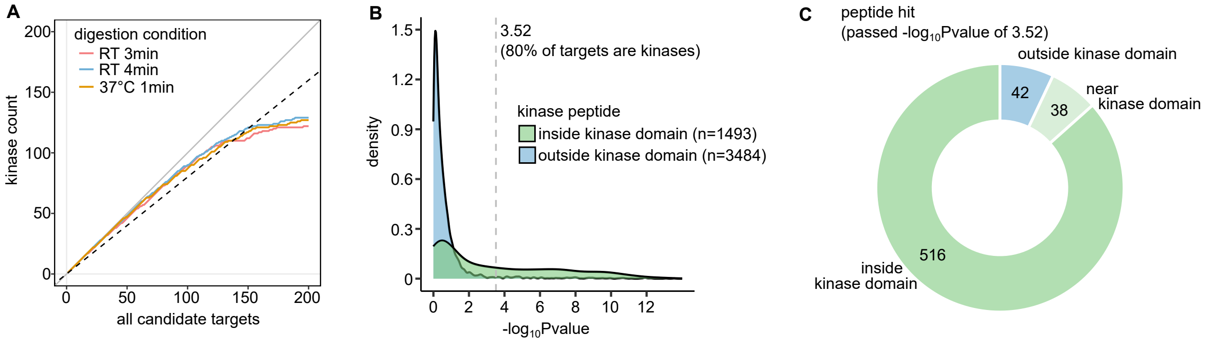

### Supplementary Figure 2

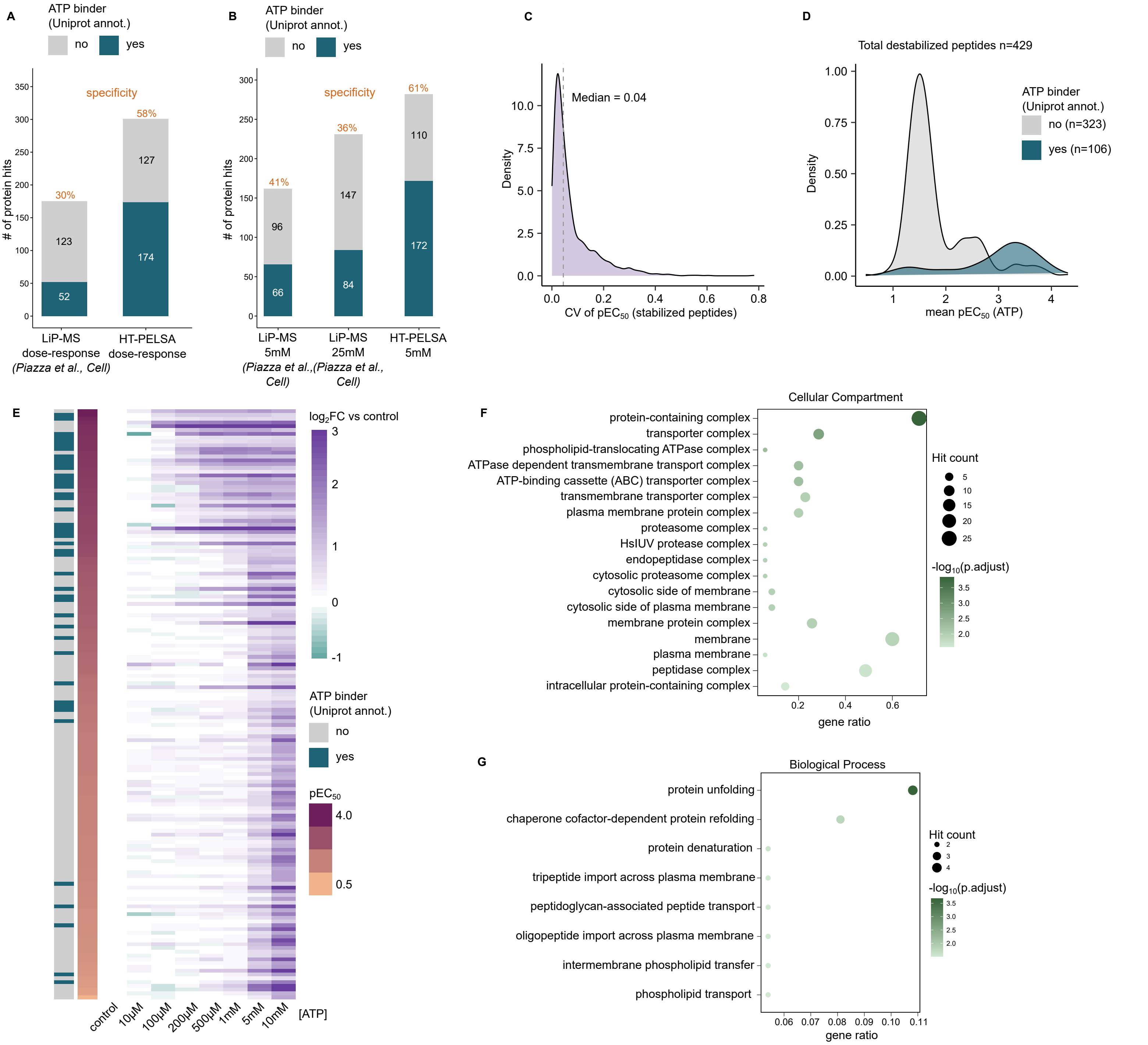

### Supplementary Figure 3

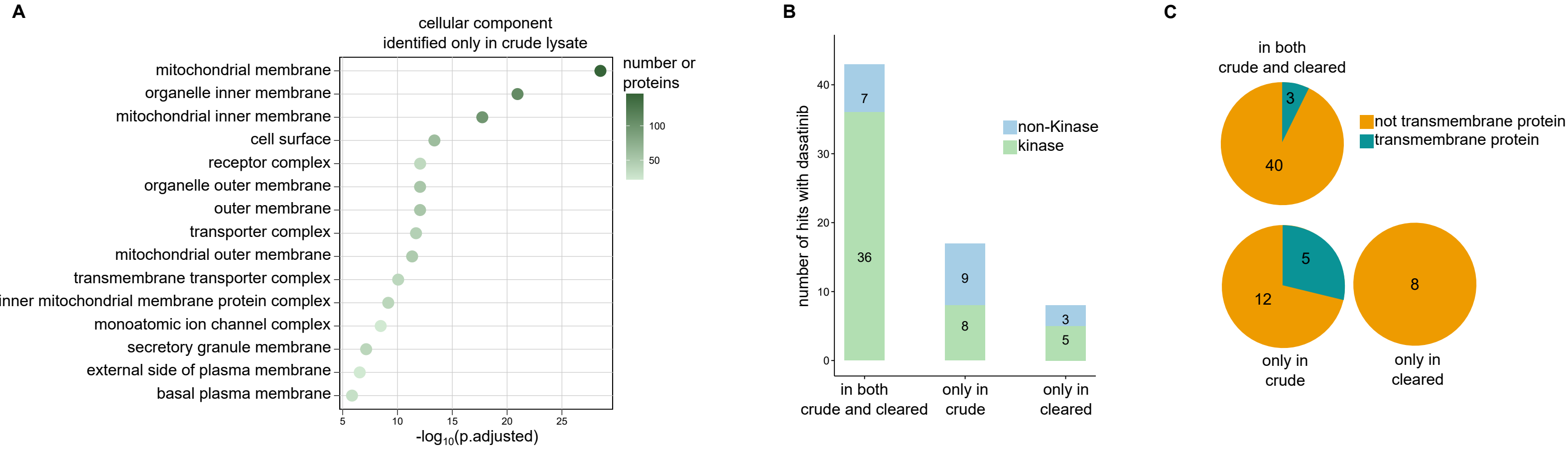
